# Auto-loaded TRAIL-exosomes derived from induced neural stem cells for brain cancer therapy

**DOI:** 10.1101/2024.05.24.595724

**Authors:** Xiaopei Zhang, Hannah Taylor, Alain Valdivia, Rajaneekar Dasari, Andrew Buckley, Emily Bonacquisti, Juliane Nguyen, Krishna Kanchi, David L. Corcoran, Laura E. Herring, Dennis A. Steindler, Albert Baldwin, Shawn Hingtgen, Andrew Benson Satterlee

## Abstract

Transdifferentiation (TD), a somatic cell reprogramming process that eliminates pluripotent intermediates, creates cells that are ideal for personalized anti-cancer therapy. Here, we provide the first evidence that extracellular vesicles (EVs) from TD-derived induced neural stem cells (Exo-iNSCs) are an efficacious treatment strategy for brain cancer. We found that genetically engineered iNSCs generated EVs loaded with the tumoricidal gene product TRAIL at nearly twice the rate as their parental fibroblasts, and the TRAIL produced by iNSCs were naturally loaded into the lumen of EVs and arrayed across their outer membrane (Exo-iNSC-TRAIL). Uptake studies in *ex vivo* organotypic brain slice cultures showed Exo-iNSC-TRAIL selectively accumulates within tumor foci, and co-culture assays showed that Exo-iNSC-TRAIL killed metastatic and primary brain cancer cells more effectively than free TRAIL. In an orthotopic mouse model of brain cancer, Exo-iNSC-TRAIL reduced breast-to-brain tumor xenografts around 3000-fold greater than treatment with free TRAIL, with all Exo-iNSC-TRAIL treated animals surviving through 90 days post-treatment. In additional *in vivo* testing against aggressive U87 and invasive GBM8 glioblastoma tumors, Exo-iNSC-TRAIL also induced a statistically significant increase in survival. These studies establish a new easily generated, stable, tumor-targeted EV to efficaciously treat multiple forms of brain cancer.

## 1. Introduction

Malignant brain tumors present diverse and significant therapeutic hurdles, and there is a desperate need for new, creative, approaches to treatment[1],[2],[3]. Standard resection and chemo-radiation treatment lack the ability to eradicate diffuse malignant cells; therefore, recurrence of some brain tumors is almost unavoidable. Over the past two decades, engineered tumoricidal neural stem cells (NSCs) have broken new ground as potential therapeutics, demonstrating promise as an alternative cancer treatment approach via targeted migration to brain tumor cells and delivery of a broad selection of therapeutic agents[4],[5],[6]. By transdifferentiating fibroblast cells directly into NSCs without a pluripotent intermediate, our lab has previously created a powerful tumor-homing and tumoricidal induced NSC (iNSC) from human skin fibroblasts[7],[8],[9]. These iNSCs have been engineered to produce and secrete a number of therapeutic agents which are actively delivered to tumor cells through a robust tumor-homing capacity. This active migration into invasive tumor foci can increase survival by delivering therapeutics into brain tumor regions that conventional surgery, chemotherapy, and radiotherapy routinely miss[7],[8],[9],[10],[11]. Despite the success of these studies, many concerns, ranging from complex manufacturing, enormous production costs, and potentially harmful side effects, remain major limitations of the clinical benefits of iNSC therapy [7],[12],[13]. One way to maintain the robust potency of cytotoxic cell therapies while mitigating regulatory, financial, and technical limitations may be to isolate and deliver only the secreted therapeutic product [14],[15],[16].

Extracellular vesicles (EVs) and exosomes are nanosized, membrane-bound vesicles that range in size from 30-200 nm [17]. They are constituted of many substances, primarily including proteins, lipids, and nucleic acids [18], and can be identified by enrichment of proteins such as tetraspanins (e.g. CD63, CD9, and CD81) and membrane-binding proteins such as Tsg101 [19]. EVs are produced intracellularly during the process of plasma membrane invagination and multivesicular body (MVB) development and are ultimately secreted via exocytosis from MVBs by fusion with the cellular plasma membrane [20],[21]. EVs are secreted by virtually all types of cells, including NSCs, and have been proposed as a potential alternative to treatment with cells.[22] EVs have been referred to as “miniature surrogates” of their parental cells, because they can partially inherit analogous therapeutic properties from their original cells [23],[24]. EVs naturally secreted from stem cells (e.g., NSCs, induced pluripotent stem cells (iPSCs), and mesenchymal stem cells (MSCs)) have diverse therapeutic properties, including anti-inflammation, immunity modulation, and tissue repair [25]. Furthermore, stem cell-derived EVs can be more easily isolated and preserved, have higher safety and immune tolerance, and present fewer ethical issues when compared to the full cell product [26],[23]. While NSC-derived EV therapeutics have been studied for treating neurodegeneration and cancer, studies on EVs derived from iNSCs for brain cancer therapy have not yet been reported [27],[28],[29].

TNF-related apoptosis-inducing ligand (TRAIL), a type-II transmembrane protein of the TNF superfamily, has long held promise as a cancer therapy [30],[31]. TRAIL targets the extrinsic apoptotic pathway, triggering caspase-induced apoptosis in malignant cells with high expression of Death Receptor 4 or 5 (DR4/5) while sparing healthy cells [32]. While TRAIL has shown strong antitumoral effect in preclinical investigations, clinical trials have shown that TRAIL alone is insufficient to effectively treat patients [33],[34]. This could be due to its poor pharmacokinetics, such as short chemical and biological half-lives, insufficient distribution to target areas, inefficiency in stimulating DR4 and DR5 receptors of tumor cells and acquired TRAIL resistance [35],[36].To help circumvent these constraints, robust TRAIL delivery is required.

Here, we report on therapeutic EVs derived from iNSCs that have been engineered to produce and secrete TRAIL. We show for the first time that a significant proportion of TRAIL is secreted by iNSCs via EVs, which are fully made, loaded, and secreted by the cells themselves. We now examine the therapeutic potential of these EVs, termed Exo-iNSC-TRAIL (**Fig. 1**), isolated from TRAIL-secreting iNSCs *in vitro.* Characterization and *in vitro/ex vivo/in vivo* testing of Exo-iNSC-TRAIL as a local brain cancer therapy demonstrate its tumor-selective accumulation and therapeutic superiority to free TRAIL protein in three orthotopic models of glioblastoma and breast cancer brain metastasis.

**Fig. 1.**
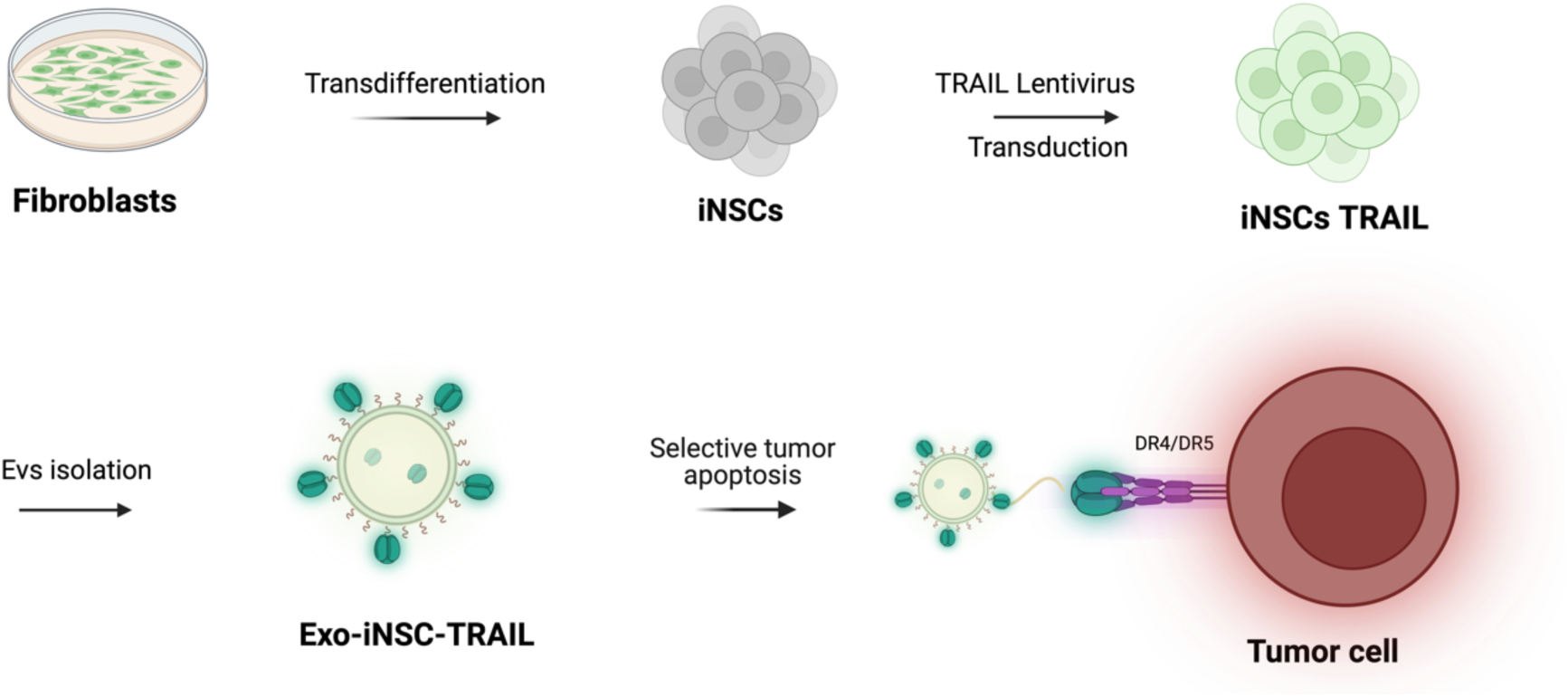
Scheme demonstrating the generation and isolation of Exo-iNSC-TRAIL from iNSCs and the selective cytotoxicity of Exo-iNSC-TRAIL to tumor cells.

## 2. Material and methods

### Materials

TRAIL was purchased from Millipore sigma (CAT No.GF092). TRAIL ELISA kit was purchased from R&D Systems (CAT NO. Invitrogen™ BMS2004). Uranyl Acetate Solution (2%) (CAT NO. 22400-2) were purchased from Electron Micros. Caspase-Glo® 3/7 Assay System (CAT NO. G8090), Caspase-Glo® 8 Assay Systems (CAT NO. G8200), and Caspase-Glo® 9 Assay Systems (CAT NO. G8210), were purchased from Promega. Pierce™ BCA Protein Assay Kit (CAT NO. 23227), RIPA Lysis and Extraction Buffer (CAT NO. 89901), and DAPI (4’,6-Diamidino-2-Phenylindole, Dilactate) (CAT NO. D3571) were purchased from Thermo Fisher Scientific. PKH67 Green Fluorescent Cell Linker Mini Kit for General Cell Membrane Labeling (CAT NO. MINI67) and anti-GFAP Cy3 conjugate (CAT NO. MAB 3402C3) were purchased from Millipore Sigma. PE anti-human CD253 (Trail) (CAT NO. 308206) was purchased from Biolegend. CD63 Monoclonal Antibody (H5C6), eFluor™ 660, (CAT NO. 50-0639-42) was purchased Thermo Fsher Scientific. Goat anti-mouse IgG H&L (6 nm Gold) (CAT NO. ab39614) and anti-TSG101 antibody (CAT NO. ab30871) were purchased from Abcam. Alexa Fluor 568 goat anti-rabbit IgG (H+L) secondary antibody (CAT NO. A-11011) was purchased from Invitrogen. Anti-IBA1 antibody was purchased from (CAT NO. 019-19741) FUJIFILM Wako Pure Chemical Corporation. Lentivirus expressing Firefly Luciferase-mCherry, puromycin selection and TRAIL-GFP-puro lentivirus non-concentrated were purchased from Duke Viral Vector Core.

### Cell lines

Cell lines were received and used as described previously [9],[8]. Briefly, the treatment of triple negative breast cancer (TNBC) brain metastasis cell line MDA-MB-231-Br was obtained through material transfer agreement (MTA) (T. Yoneda). MDA-MB231-Br were cultured in Dulbecco’s modified Eagle’s medium (DMEM) supplemented with 10% fetal bovine serum (FBS) (v/v), 1% penicillin-streptomycin (p/s)(v/v). U87 glioma cells were purchased from the American Type Culture Collection (ATCC) and cultured MEM with 10% FBS and 1% p/s. Human fibroblasts cells (NHF1) were provided by W. Kauffman (The University of North Carolina School of Medicine) and were cultured in DMEM with 10% FBS and 1% p/s. GBM8 glioma neurospheres were a gift from H. Wakimoto (Massachusetts General Hospital) and were cultured in EF medium (neurobasal media, 1% L-glutamine, 1× B27 supplement, 0.5× N-2 supplement, 2 μg/mL heparin, 20 ng/mL recombinant human epidermal growth factor, 20 ng/mL recombinant human fibroblast growth factor 2). iNSCs spheres were generated as described previously and maintained in ReNcell NSC Maintenance Media [9]. Lentiviral infection of cell lines was completed as described previously [37].

### Animal ethics statement

The Institutional Animal Care and Use Committee at the University of North Carolina-Chapel Hill approved all work performed on female athymic nude mice (therapy investigation) and Sprague-Dawley rats (OBSC preparation). As described previously, all OBSCs originate from P8 Sprague-Dawley rat pups. Sprague-Dawley rats are housed with a mother and ten pups at a time. Pups are used before having to be weaned. All athymic nude mice are female and reside in cages of five or fewer [1].

### Isolation and characterization of EVs

The iNSCs or NHF1 cells were cultured in EV-depleted ReNcell NSC maintenance and DMEM medium at 37 °C respectively, and the conditioned media (CM) was collected after 48 h. The CM was then centrifuged at 1500 rpm for 15 min and 12,000 g for 20 min to get rid of cells and debris. EVs were isolated by ultracentrifugation of the supernatants at 100,000g for 70 min at 4 °C. EVs were then washed in PBS at 100,000 g for 70 min at 4 °C [38]. The isolated EVs were then reconstituted with PBS and stored at 4°C. EV amounts were quantified via protein concentration by a BCA protein assay kit.

### Transmission electron microscope and Immunoelectronmicroscopy

The morphology of EVs was observed by Transmission Electron Microscope (TEM) via uranyl acetate negative staining [39]. To observe the EV surface TRAIL, immuno-electron microscopy was applied to stain and observe the TRAIL. Briefly, EVs were fixed with 4% paraformaldehyde and dropped on carbon-coated copper grids. TRAIL antibodies (1:5) were co-incubated with fixed EVs at 4 °C overnight, and 6 nm gold-conjugated goat anti-mouse IgG secondary antibodies were then used to label the conjugated TRAIL [40]. EVs were observed by TEM from Thermo Scientific™ Talos™ F200X.

### ELISA

The level of TRAIL expressed in intact and lysed iNSC-derived EVs was quantified using ELISA following the directions provided by the manufacturer. Lysing buffer was supplemented with a protease inhibitor which did not affect TRAIL stability, as shown in Fig S1.

### Flow cytometry for Exo-iNSC-TRAIL

Approximately 400 μg/ml of isolated Exo-iNSC-TRAIL were stained with CD63 eFluor 660 and TRAIL-PE at a ratio of 1:100 (v/v) overnight at 4 °C. The unbounded antibodies were removed by ultracentrifuge at 100000 g for 70 min and washed one time with PBS. Flow cytometry for Exo-iNSC-TRAIL was performed on a ImageStreamX Mark II flow cytometry.

### Proteomics analysis

Protein exosome samples (n = 3) were diluted with 100 µL of 8 M urea in 50mM ammonium bicarbonate, pH 7.8. Samples were then reduced with 100 mM DTT at 37 ⁰C for 45 min and alkylated with 15 mM iodoacetamide for 45 min at room temperature. Samples were then diluted to 1 M urea and subjected to digestion with trypsin (Promega, 1:50 w/w) overnight at 37⁰C. The resulting peptides were acidified to 0.5 % trifluoroacetic acid, cleaned using desalting spin columns (Thermo), and eluates were dried via vacuum centrifugation. The resultant peptide samples were quantified by Pierce Fluorometric Peptide Assay, then normalized to 0.4 µg/µL prior to liquid chromatography-tandem mass spectrometry (LC-MS/MS) analysis.

Samples were analyzed by LC-MS/MS using an Easy nLC 1200 coupled to a QExactive HF mass spectrometer (Thermo Scientific). Samples were injected onto an Easy Spray PepMap C18 column (75 μm id × 25 cm, 2 μm particle size) (Thermo Scientific) and separated over a 120 min method. The gradient for separation consisted of 5–45% mobile phase B at a 250 nl/min flow rate, where mobile phase A was 0.1% formic acid in water and mobile phase B consisted of 0.1% formic acid in 80% ACN. The QExactive HF was operated in data-dependent mode (DDA) where the 15 most intense precursors were selected for subsequent fragmentation. Resolution for the precursor scan (m/z 350–1700) was set to 60,000, while MS/MS scans resolution was set to 15,000. The normalized collision energy was set to 27% for HCD. Peptide match was set to preferred, and precursors with unknown charge or a charge state of 1 and ≥ 7 were excluded.

Raw data were processed using Proteome Discoverer (Thermo Scientific, version 2.5). Data were searched against a reviewed Uniprot human database (containing ∼20,000 sequences), appended with a common contaminants database, using the Sequest HT search algorithm within Proteome Discoverer. Enzyme specificity was set to trypsin, up to two missed cleavage sites were allowed, carbamidomethylation of Cys was set as fix modification and oxidation of Met was set as variable modification. A precursor mass tolerance of 10ppm and fragment mass tolerance of 0.02 Da were used. Label-free quantification (LFQ) using razor + unique peptides was enabled through the Minora node. A 1% peptide false discovery rate (FDR), and a minimum of 2 peptides were used to filter the data. Proteins with > 3 missing LFQ intensities across samples were removed. Further analysis (log2 transformation, normalization, imputation, statistical analysis) was conducted using Argonaut [41]. Differentially abundant proteins with a FDR adjusted p-value < 0.01 were analyzed via Reactome Pathway Browser to test for pathway enrichment and pathway topology analysis [42]. The false discovery rate was used to correct for multiple hypothesis testing.

### iNSC immunohistochemistry staining

iNSC-TRAIL were seeded on the laminin coated 15 mm glass cover and cultured for 24h. iNSCs were then fixed with 10% neutral buffered formalin and permeabilized in 0.1% Triton X-100 in phosphate buffered saline (PBS-T) with 1% (w/v) bovine serum albumin (BSA). iNSCs were then incubated with mouse anti human TRAIL-PE conjugate (1:1000, v/v), mouse anti human CD63 eFluor 660 conjugate (1:1000, v/v), and primary rabbit anti human TSG101 (1:1000, v/v) antibodies for 2 h at room temperature. For TSG fluorescence labeled secondary antibody, iNSCs were washed with PBS three times and then incubated with Alexa Fluor 568 goat anti-rabbit (1:1000, v/v) secondary antibodies for 1 h. Then iNSCs were washed with PBS three times and stained with nuclei DAPI stain.[43] After that, iNSCs were mounted on the slides by Pro-Long Gold Antifade Mountant. The slides were observed by Leica SPX8 confocal microscopy. Representative images were analyzed by Leica image software.

### Post-loading of free TRAIL onto Exo-iNSC

Blank Exo-iNSCs were isolated from iNSCs that did not express TRAIL, while free TRAIL was subsequently isolated from iNSC-TRAIL by removing Exo-iNSC-TRAIL via ultracentrifuge. Either 1.5 ng or 3 ng of the isolated free TRAIL (iNSC) was incubated with 1 μg of the blank EVs derived from iNSC overnight, and the EVs were subsequently washed to remove the free TRAIL. The TRAIL associated with Exo-iNSC was measured by TRAIL ELISA kit.

### In vitro studies

100 μL of MDA-MB231-Br-mCherry-FLuc cells (5000 cells/well), GBM8-mCherry-FLuc cells (10000 cells/well) and U87-mCherry-FLuc cells (10000 cells/well) were implanted in black 96-well clear-bottom plates. After 24h, the cell media was exchanged by fresh media and different concentrations of free TRAIL and the same amount of TRAIL from Exo-iNSC-TRAIL were added simultaneously. After co-incubation for 72h, cell viability was assessed via bioluminescence (BLI) via AMI optical imaging system. One outlying data point in Fig. 3C was removed using Tukey’s Inter-Quartile Range method.

### EVs uptake by Organotypic Brain Slice Cultures (OBSCs)

All slicing-related activities were approved by the Institutional Animal Care and Use Committee at the University of North Carolina-Chapel Hill. OBSCs were generated as described previously.[1] Briefly, OBSCs were sliced from P8 Sprague-Dawley rat pups. Dissected brains were fixed on a vibratome platform (Leica VT1000S) and immersed in ice-cold brain slice media, also defined previously.[1] Coronal OBSCs were sliced at a thickness of 300 μm. OBSC were then cultured on 6-well matched Millicell culture inserts with 1 mL of brain slice media underneath each insert. For tumor implantation on OBSCs,1 μL of 20,000 MDA-MB231-Br-mCherry-FLuc cells were added on the center of each OBSC hemisphere at 2 h after OBSC generation. Fresh OBSC medium was changed the day after slicing, and 100 μg/mL PKH26-labeled EVs were added into the OBSC media. After 24 h, the OBSCs were washed with PBS and fixed with 4% paraformaldehyde for immunofluorescent staining.

### Immunofluorescence staining of OBSCs

Fixed OBSCs were first washed by PBS three times. 0.1% Triton X-100 in phosphate buffered saline (PBS-T) was then used to permeabilize the OBSCs for 1 h. OBSCs were then blocked in PBST with 1% BSA for 1h at room temperature. OBSCs were incubated with Cy3 conjugated glial fibrillary acidic protein (GFAP,1:1000 v/v), and ionized calcium-binding adapter molecule 1 (IBA1, 1:1000 v/v) overnight. After washed by PBS three times, OBSCs were stained with secondary antibody Alexa Fluor 568 goat anti-rabbit IgG (1:1000 v/v) 1 h. Nuclei were stained by DAPI [44]. Finally, OBSCs were mounted on the slides by Pro-Long Gold Antifade liquid mountant and observed by Leica SPX8 confocal microscopy at UNC Neuroscience Microscopy Core.

### In vivo BLI

To monitor the tumor growth of MDA-MB231-Br-mCherry-FLuc cells, GBM8-mCherry-FLuc, and U87-mCherry-FLuc, sequential BLI was conducted by the following method. Each mouse was intraperitoneally injected with 200 μL of 1.5 mg/ml d-luciferin in PBS. 10 min later, BLI was performed on the luciferin-administered mice by an AMI optical imager system, as described previously [9],[11]. BLI data were analyzed with Aura software.

### IC-infused therapy studies in TNBC-brain metastasis model

MDA-MB231-Br-mCherry-FLuc cells (8 × 10^4^) in 3 μL of PBS were implanted in the right hemisphere of the brain in athymic nude mice by stereotactic intracranial (IC) injection, as described previously.[11] Four days after tumor implantation, a single dose of 8 ng free TRAIL or an equivalent TRAIL dose of Exo-iNSC-TRAIL in 6 μL of PBS was infused into the tumors. For the control group, 6 μL of PBS was injected in the brain by the same protocol. Serial BLI was completed via the AMI optical imager system to monitor the growth of tumors in mice. When a mouse’s body weight decreased by over 20% of its original weight, it was sacrificed, and data on survival was recorded.

### IC-infused therapy studies in Glioblastoma models

GBM8-mCherry-FLuc cells (1 × 10^5^) and U87-mCherry-FLuc cells (1 × 10^5^) in 3 μL of PBS were implanted in the right hemisphere of the brain in athymic nude mice by stereotactic intracranial (IC) injection as described previously.[11] For the GBM8 model, 8 ng free TRAIL or an equivalent TRAIL dose of Exo-iNSC-TRAIL in 6 μL of PBS was injected IC at day 4, 11 and 25 after tumor implantation. For the U87 model, a single dose of 8 ng free TRAIL or an equivalent TRAIL dose of Exo-iNSC-TRAIL in 6 μL of PBS was infused into the tumors on day 4 after tumor implantation. For the control group, 6 μL of PBS was injected in the brain by the same protocol. Serial BLI was completed via the AMI optical imaging system to monitor the growth of tumors in mice. When a mouse’s body weight decreased by over 20% of its original weight, it was sacrificed, and data on survival was recorded.

## 3. Results

We recently developed a second-generation TRAIL-overexpressing iNSC cell type that exhibits improved antitumor properties and significantly increased tumor-homing capability.[9] Since TRAIL protein can be expressed in both membranous and secreted formats, we hypothesized that TRAIL was being released from iNSCs not only as free protein, but also associated with iNSC-derived EVs. To investigate this, TRAIL and canonical EVs markers CD63 and TS101 were first intracellularly co-stained in iNSCs (**Fig. 2A**). Represented fluorescence images reveal the co-localization of TRAIL and EVs within the iNSCs, indicating that TRAIL may be loaded into EVs following EVs secretion. EVs were isolated from conditioned iNSC culture media using a serial centrifugation procedure.[45] The successfully isolated EVs were first confirmed by transmission electron microscopy (TEM) ( **Fig. 2B**). The represented TEM images show these isolated particles exhibit typical EVs characteristics including a cup-like morphology and a diameter of 50 to 200 nm. Dynamic Light Scattering (DLS) and Nanoparticle Tracking Analysis (NTA) confirmed a consistent nanoparticle population with a zeta-potential of - 24 ± 3 and an average size of 138 ± 7 nm (**Fig. 2C**). The isolated particles were also labeled with CD63 to further validate their EVs property. Flow cytometry of CD63 labeled particles revealed that ∼80% of isolated particles were CD63-positive, and these populations maintained a similar size distribution to the unlabeled population, indicating that CD63-containing EVs had indeed been isolated and termed as Exo-iNSC (**Supplementary Fig.S2**). Interestingly, the proteome encapsulated within Exo-iNSC-TRAIL was significantly different than the proteome in EVs derived from pre-transdifferentiated NHF1 fibroblasts (Exo-NHF1-TRAIL) (p<0.01, n = 3 biological replicates, **Supplementary Fig. S3**).

**Fig. 2.**
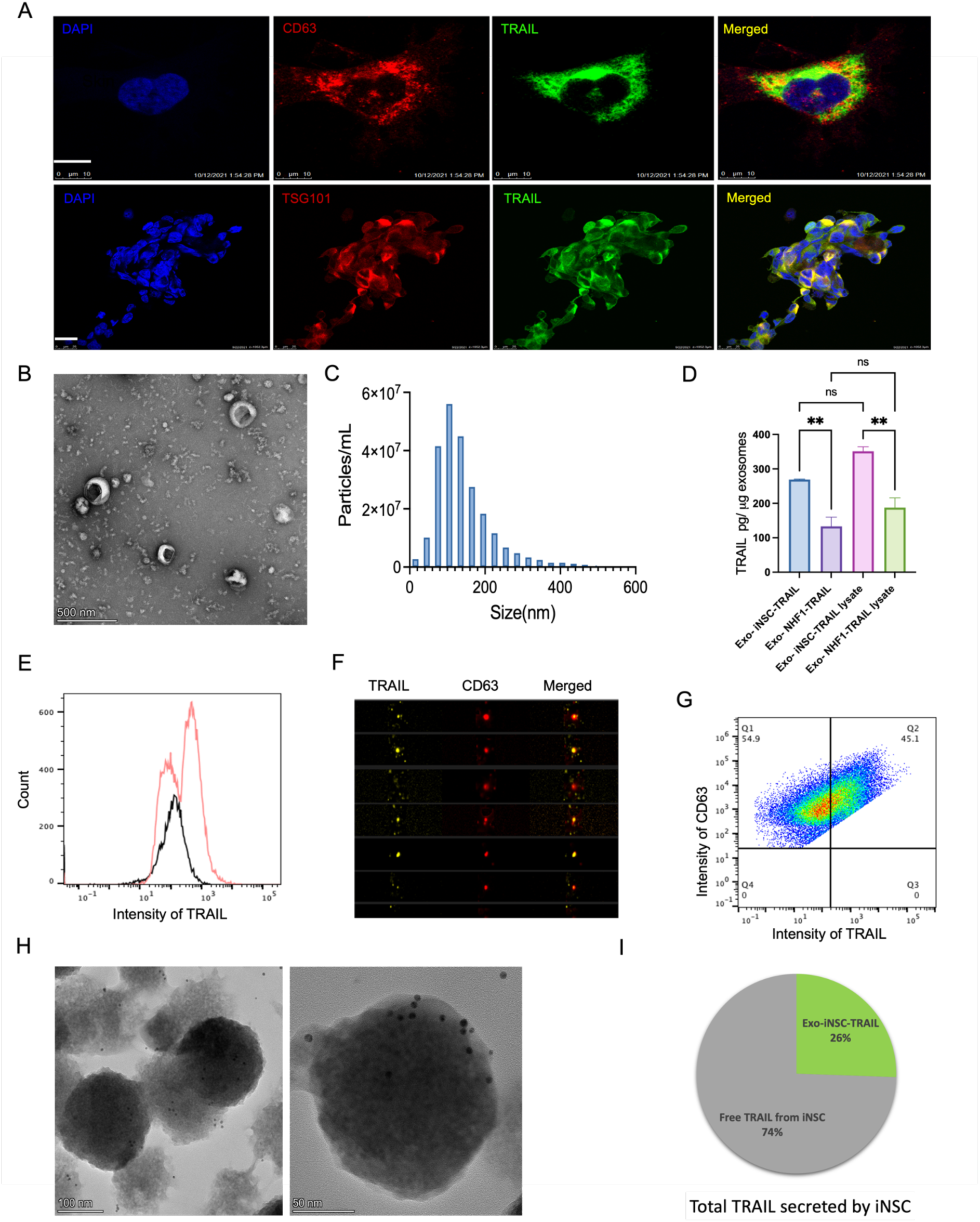
Isolation, characterization and TRAIL measurement of Exo-iNSC-TRAIL. **A**, Co-localization of TRAIL and exosomes inside iNSCs. Upper, representative confocal microscopy images of TRAIL-exosome colocalization in a single cell. Lower, representative confocal microscopy images of TRAIL-exosome colocalization in iNSC spheres. Exosomes were stained by CD63 (Red, upper) and TSG101 (Red, lower), TRAIL was stained by TRAIL antibody (green), and nuclei of iNSC spheres were labelled with DAPI (blue). Scale bar represents 10 µm (upper) and 25 µm (lower); **B**, negative contrast micrograph of Exo-iNSC-TRAIL examined and imaged by transmission electron microscopy (TEM), scale bar 500 nm; **C**, NTA characterizes the size distribution of Exo-iNSC-TRAIL; **D**, the measurement of TRAIL expressed on intact Exo-iNSC-TRAIL, intact Exo-NHF1-TRAIL, Exo-iNSC-TRAIL lysate, and Exo-NHF1-TRAIL lysate by ELISA, analyzed by one-way ANOVA (**P < 0.01); **E**, flow cytometry detected the percentage of TRAIL expression on the iNSC Exo-TRAIL surface; **F-G**, TRAIL expression on Exo-iNSC-TRAIL confirmed by dual-labelling of exosomes for TRAIL and CD63 and quantification/imaging via flow cytometry; **H**, representative immuno-electron-microscopy images of Exo-iNSC-TRAIL stained with TRAIL primary antibody and 6-nm nanogold secondary antibody; **I**, the percentage of the total TRAIL secreted by iNSC-TRAIL cells that is associated with Exo-iNSC-TRAIL (n=6 replicates). All data are shown as mean ± SEM.

**Fig. 3.**
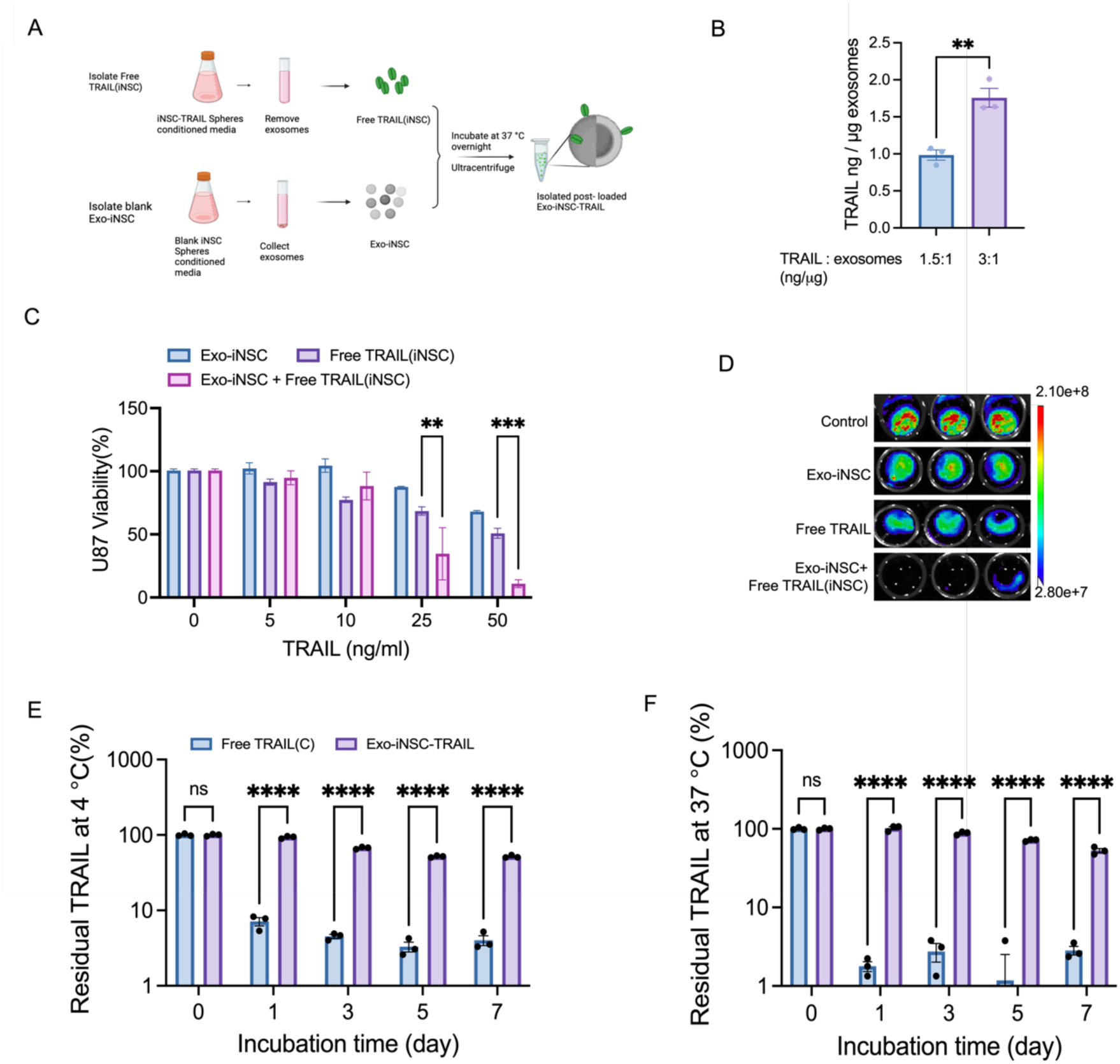
Toxicity and stability of TRAIL is higher after associated with surface of Exo-iNSC. **A**, The scheme to demonstrate post-loading of free TRAIL onto Exo-iNSC; **B**, Free TRAIL associates with Exo-iNSC after co-incubation and washing (measured by ELISA), analyzed by t-test (\*\**P* <0.01); **C**, post-loaded Exo-iNSC-TRAIL displays enhanced cytotoxicity compared to free TRAIL, analyzed by two-way ANOVA (\*\**P* <0.01 and \*\*\**P* < 0.001); **D**, the representative BLI images of U87 viability after treatment with 25 ng/mL TRAIL for 48h, analyzed two-way ANOVA (n = 4-5, \*\*\*\**P* < 0.0001);**E** and **F**, the thermal stability of the TRAIL in Exo-iNSC-TRAIL after incubation at 4 ℃ and 37 ℃ for different time intervals, analyzed by two-way ANOVA (n = 3, \*\*\*\**P* < 0.0001). All data are shown as mean ± SEM.

Next, we looked into whether the iNSC-secreted EVs contained functional TRAIL. A highly specific commercial ELISA kit was used to investigate the expression of TRAIL in prepared iNSC EVs. Total TRAIL expression in Exo-iNSC-TRAIL was higher than TRAIL expression in EVs generated from pre-transdifferentiated NHF1 fibroblasts (**Fig. 2D**). Exo-iNSC-TRAIL reproducibly contained nearly double the TRAIL per EV than Exo-NHF1-TRAIL (351±13.3 pg TRAIL/ug EVs vs ∼187 ± 28.4 pg TRAIL/ug EVs). Interestingly, ELISA quantification of TRAIL on the surface of intact Exo-iNSC-TRAIL (∼269.5 ± 1.4 pg TRAIL/ug EVs) was similar to the total EV-associated TRAIL quantified from Exo-iNSC-TRAIL lysate (∼351±13.3 pg TRAIL/ug EVs) (**Fig. 2D**). Lysing buffer was supplemented with a protease inhibitor which did not affect TRAIL stability, as shown in Fig S1. This suggested that TRAIL associated with Exo-iNSC-TRAIL was largely bound to the EV surface (∼75% of total TRAIL). TRAIL arrayed on the surface of EVs was also validated by image flow cytometry by using the TRAIL-specific and CD63-specific antibodies. Results confirmed that ∼42% of the isolated vesicles were TRAIL positive and ∼45% of CD63+ derived EVs expressed with TRAIL (**Fig. 2E-G**). Moreover, TRAIL expression on the surface of Exo-iNSC-TRAIL was examined by nanogold immunostaining (**Fig. 2H**). Immuno-electron-microscopy analysis showed vesicles highly positive for surface TRAIL, consistent with our previous results. Using a commercial ELISA, TRAIL expression on Exo-iNSC-TRAIL was compared to the total TRAIL secreted by iNSCs, revealing approximately 26% of the TRAIL produced by iNSCs was associated with Exo-iNSC-TRAIL (**Fig. 2I**). Taken together, these data suggest that iNSCs produce and secrete a significant percent of TRAIL within a uniform population of EVs, and that TRAIL was not only loaded into the lumen of the EVs but distributed across the membrane of the particles as well.

To determine whether TRAIL is attached to EV surfaces before or after secretion, we designed an experiment to determine whether TRAIL can conjugate to surfaces of EVs after EV isolation (**Fig. 3A**). Blank Exo-iNSCs were isolated from iNSCs that did not express TRAIL, while free TRAIL was subsequently isolated from iNSC-TRAIL, described as free TRAIL (iNSC). Either 1.5 or 3 ng of the isolated free TRAIL (iNSC) was incubated with 1 μg of the blank EVs overnight, and the EVs were subsequently washed to remove the free TRAIL. ELISA measured a very large amount of TRAIL associated with Exo-iNSC (∼1000 pg TRAIL/ μg EVs and ∼1758 pg TRAIL/ μg EVs respectively, **Fig 3B**), suggesting that free TRAIL can naturally associate with constituents on the outer membrane of Exo-iNSC. It’s worth noting that EVs derived from non-TRAIL-expressing iNSCs were negative for TRAIL expression. Interestingly, “post-loaded” Exo-iNSC-TRAIL exhibited significantly greater tumor kill against the glioblastoma cell line U87 *in vitro* when compared to the same dose of free TRAIL (**Fig. 3C-D**). At a dose of 25 ng/mL, Exo-iNSC-TRAIL induced ∼85% kill at t = 48 h after treatment compared to just ∼30% kill by free TRAIL (iNSC). This could be due to the greater stability of TRAIL when associated with EVs (**Fig. 3E-F**).

One of the issues preventing the production of recombinant free TRAIL agent is their rapid degradation and clearance when exposed to harsh conditions such as the fluctuating temperature during transportation. Studies have shown that EVs can stabilize TRAIL proteins which could augment their anti-cancer effects. Here we show that the stability of TRAIL within Exo-iNSC-TRAIL is significantly enhanced. The stability of the free TRAIL and Exo-iNSC-TRAIL was investigated by monitoring the TRAIL concentration after incubation at 4°C and 37 °C for different time intervals (**Fig. 3E-F**). The results show that incubation of free TRAIL at 4°C and 37°C caused above 90% degradation of the free protein within 1 day. In contrast, the levels of TRAIL in EVs were reduced by only ∼ 10% at 4°C and 37°C after 3 days and ∼50% after a week, suggesting the EV thermally stabilizes TRAIL and should allow prolonged activity that could enhance anti-tumor efficacy.(**Fig. 3E-F**).

The proapoptotic potential of Exo-iNSC-TRAIL was next investigated in vitro against the breast cancer brain metastasis cell line MB231Br, the glioma cell line U87, and primary patient-derived stem-like GBM8 cells. According to previous reports, MB231Br were susceptible to TRAIL and had an IC50 of approximately 2 ng/mL, whereas U87 and GBM8 cells were more resistant to TRAIL-induced apotosis[46],[47]. As shown in **Fig. 4A**, MB231Br shows high sensitivity to both Exo-iNSC-TRAIL and free TRAIL, with an IC50 of ∼3 ng/mL at 72 h. U87 and GBM8 cells were more resistant to free TRAIL, with GBM8 neurospheres showing no apoptosis after being treated with various concentrations of TRAIL for 72 h. Exo-iNSC-TRAIL was more effective than free TRAIL at inducing apoptosis in all three cell lines. Against U87 cells, Exo-iNSC-TRAIL induced an IC50 of 9 ng/mL, which is 5-fold lower than that of free TRAIL. Against GBM8 cells, Exo-iNSC-TRAIL induced nearly 50% killing at 100 ng/mL (**Fig. 4A**). These data suggest that the cytotoxic potency of TRAIL is increased when loaded on Exo-iNSC-TRAIL.

**Fig. 4.**
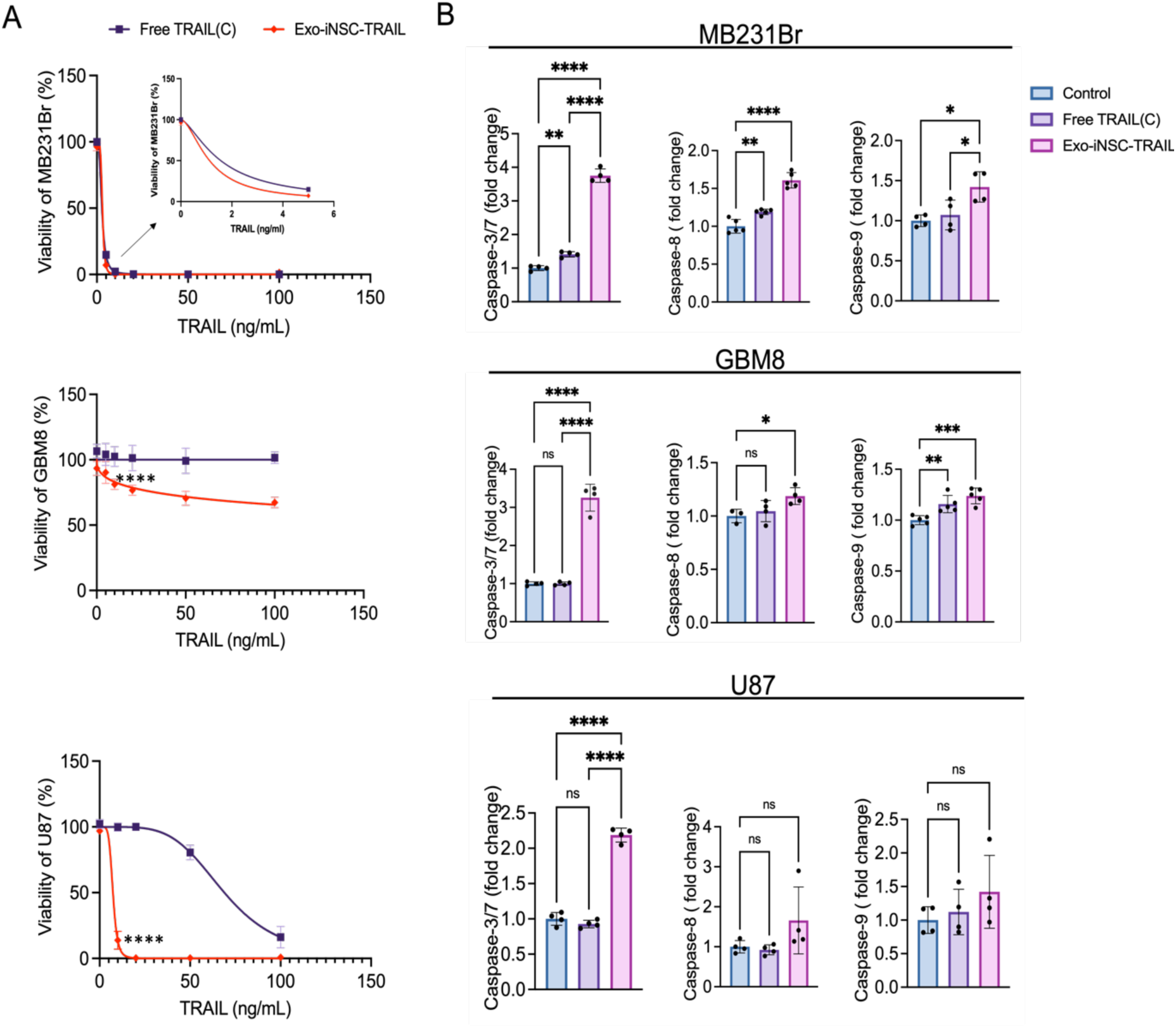
The cytotoxicity and apoptotic activity of Exo-iNSC-TRAIL on brain tumor cell lines. **A**, the cytotoxicity of Exo-iNSC-TRAIL and free TRAIL on MB231Br, GBM8, and U87 after treatment with increasing concentrations of TRAIL, t = 72h, analyzed by two-way ANOVA (\*\*\*\**P*< 0.0001); **B**, the fold change of caspase-3/7, caspase-8, and caspase-9 in MB231Br, GBM8, and U87 after treatment with 5 ng/ml of free TRAIL or Exo-iNSC-TRAIL, t = 24h, analyzed by one-way ANOVA ( \**P* < 0.05, \*\**P* < 0.01, \*\*\**P* < 0.001, and \*\*\*\**P*< 0.0001). All data are shown as mean ± SEM.

Soluble or transmembrane TRAIL can initiate the extrinsic apoptotic pathway by binding to DR4/ DR5 receptors overexpressed on many cancer cells, triggering a proteolytic cascade of caspase activation that subsequently induces apoptosis[48],[49]. To confirm that the reduction in cell viability in response to Exo-iNSC-TRAIL is mediated by apoptosis, the activation of caspase-3/7, 8 and 9 was detected MB231Br, GBM8, and U87 cell lines were treated with a low dose of TRAIL (5 ng/mL) for 24h. Very low-to-no caspase 3/7,8 and 9 activations were observed in all cells treated with free TRAIL. In contrast, around a threefold increase in caspase 3/7 activity, and around 1.4-fold increases in caspase 8 and 9 activities were observed in all 3 cell lines after treatment with Exo-iNSC-TRAIL (**Fig. 4B**). These results show that the increased potency of Exo-iNSC-TRAIL compared to free TRAIL is due to greater upregulation of apoptotic signaling rapidly after treatment initiation.

Tumor-tropic targeting of iNSC-derived EVs would be an important advantage to their potential as drug delivery vehicles. Exo-iNSC-TRAIL are derived from second-generation iNSCs that exhibit significantly increased tumor-homing capability; therefore, we hypothesized that Exo-iNSC-TRAIL may themselves have an ability to selectively target tumor cells. Using previously validated living *ex vivo* organotypic brain slice cultures (OBSCs),[1],[11],[50] we found that EVs robustly and selectively accumulate in engrafted MB231Br tumor foci while sparing normal brain tissue (**Fig. 5 and supplementary Fig. S4**). In six-well plates, OBSCs were cultured atop transwell inserts with 0.4-µm pore size and EVs were diluted in media underneath the inserts, where they were allowed to passively diffuse into OBSCs for 24h. Quantitative imaging of living OBSCs showed that iNSC-derived EVs did not accumulate at significant levels in normal brain or within activated microglia or astrocytes **(supplementary Fig. S4).** In contrast, nearly 20-fold greater levels of EVs were detected in OBSCs that had been engrafted with brain tumor foci. Throughout z-stacks of 19 10 μm-thick confocal images of OBSCs harboring MB231Br tumors, both Exo-iNSC-TRAIL and Exo-NHF1-TRAIL showed significant accumulation (**Fig. 5A**). Interestingly, while the absolute difference in EV signal between these groups was only a 1.4-fold increase in Exo-iNSC-TRAIL compared with that of Exo-NHF1-TRAIL (**Fig. 5B**), there was a significantly greater co-localization of Exo-iNSC-TRAIL within the tumor cells themselves compared to Exo-NHF1-TRAIL. An analysis of Exo/Tumor signal in each region along the z-axis revealed an ∼4-fold increase in tumor-specific accumulation of Exo-iNSC-TRAIL over Exo-NHF1-TRAIL (**Fig. 5C**). There are several possible mechanisms for this phenomenon, including (1) that Exo-iNSC-TRAIL has tropism for tumor cells, possibly via an abundance of surface-conjugated TRAIL acting as a tumor-selective targeting ligand and leading to preferential uptake over other cell types; or (2) that other, yet uncharacterized, proteins upregulated in iNSCs are transferred to Exo-iNSC-TRAIL and drive their tumor-specific accumulation.

**Fig. 5.**
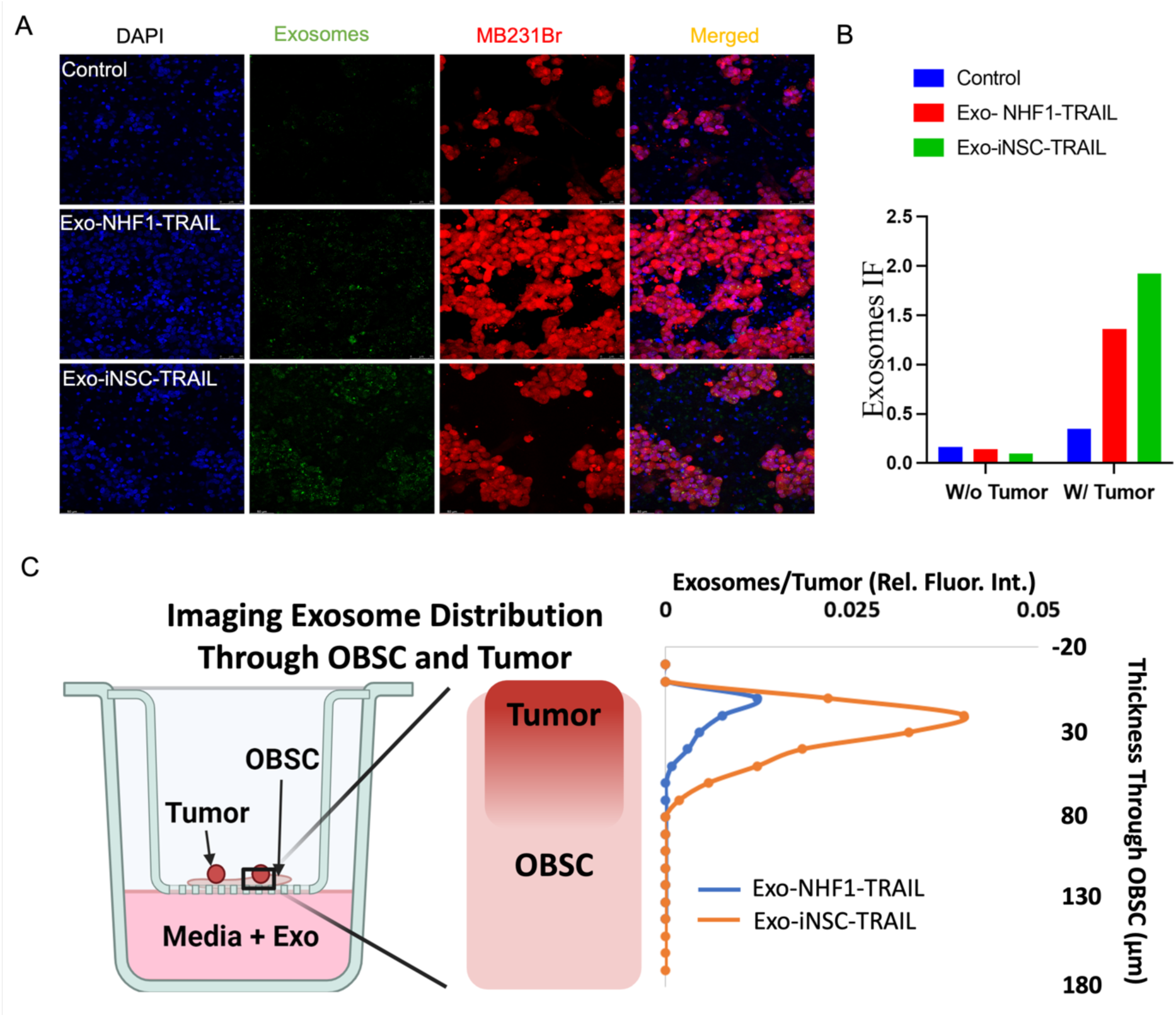
Exo-iNSC-TRAIL accumulates in tumor cells in an organotypic brain slice culture (OBSC) *ex vivo* model. **A**, representative confocal images of fluorescently labelled Exo-NHF1-TRAIL and Exo-iNSC-TRAIL in tumor-bearing OBSCs with MB231Br cells (red) engrafted atop the OBSCs. Exo-NHF1-TRAIL and Exo-iNSC-TRAIL were labelled by PKH26 (green), MB231Br were transfected with mCherry (red) and nuclei were stained by DAPI (blue). t = 24h, scale bar = 50 µM. **B**, quantification of Exo-NHF1-TRAIL and Exo-iNSC-TRAIL accumulation on OBSCs with or without tumor foci. Each bar represents the sum of 19 z-stacked images; **C**, quantification of exosome accumulation along a z-stack of 19 images which shows that Exo-iNSC-TRAIL accumulates more specifically in MB231Br tumor cells than Exo-NHF1-TRAIL cells, t = 24h.

After showing that Exo-iNSC-TRAIL were able to accumulate in the tumor foci on living brain tissue, we then proposed to explore the tumor killing potential of Exo-iNSC-TRAIL against breast-to-brain metastasis tumors in an orthotopic xenograft model of MB231Br. MB231Br cells expressing firefly luciferase were stereotactically implanted into the brain parenchyma of nude mice according to a previously validated protocol.[9] Four days after tumor implantation, a single dose of phosphate-buffered saline, 6 ng free TRAIL, or an equal amount of TRAIL in Exo-iNSC-TRAIL was directly injected into the tumor. Longitudinal bioluminescence imaging (**Fig. 6A**) revealed robust, sustained tumor killing by Exo-iNSC-TRAIL (**Fig. 6B**). In contrast, free TRAIL initially induced modest tumor growth suppression, but tumors quickly grew and many mice developed spinal tumors in addition to the primary brain mass. By 32 days after treatment, there was a statistically significant difference in brain tumor burden among animals given PBS (∼971.5-fold tumor growth), those given free TRAIL (∼1160-fold tumor growth), and Exo-iNSC-TRAIL (∼2.7-fold tumor growth) (**Fig. 6B**). While a majority of untreated animals and animals given free TRAIL had succumbed to tumor burden by 70 days after treatment, all animals who received Exo-iNSC-TRAIL showed robust and sustained tumor growth suppression (**Fig. 6C**) and survived over 90 days after treatment (**Fig. 6D**).

**Fig. 6.**
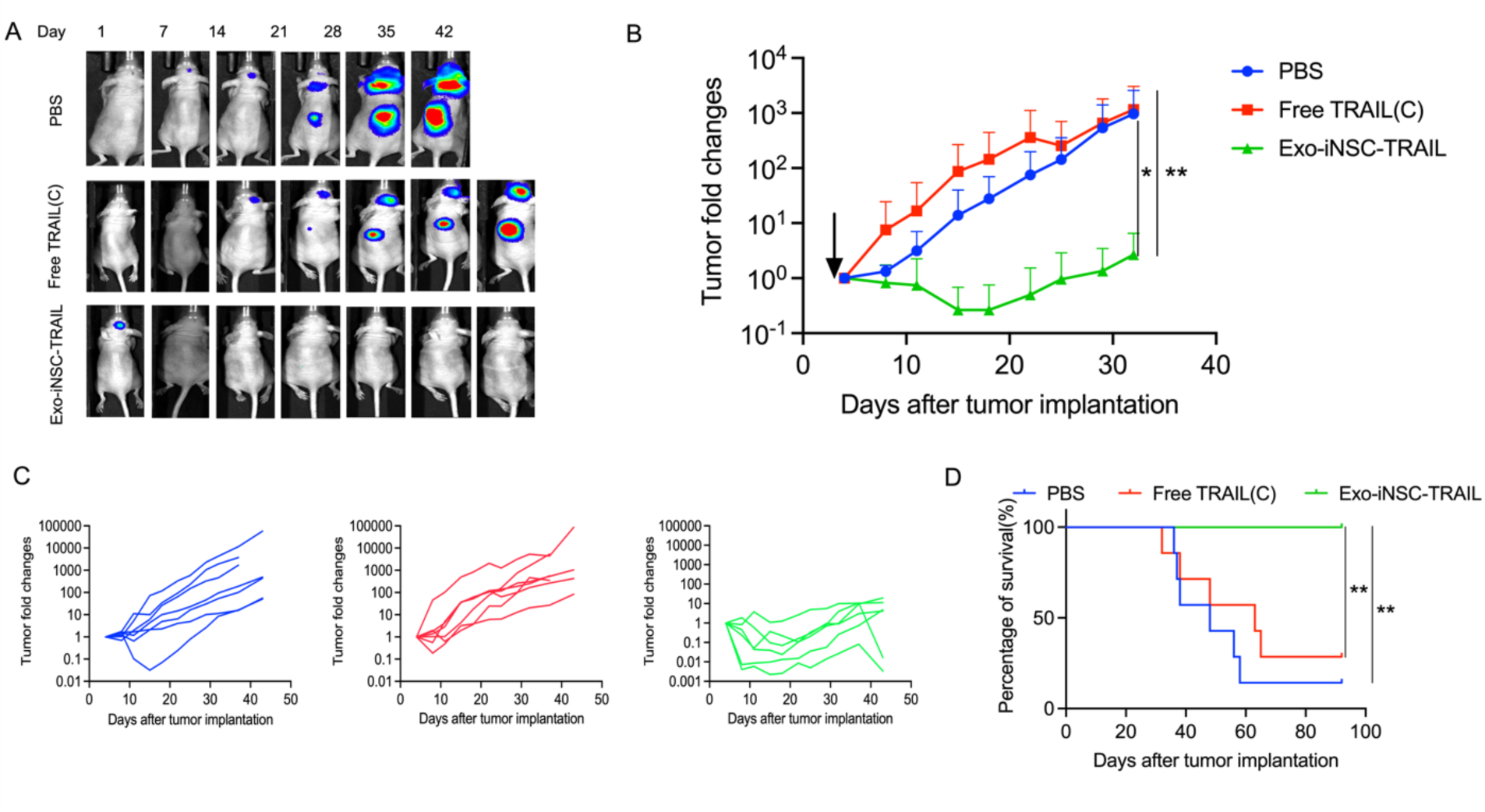
*Ex vivo* and orthotopic *in vivo* tumor killing capabilities of Exo-iNSC-TRAIL. **A**, representative BLI images of orthotopic MB231Br brain tumor-bearing mice at different time points after treatment with PBS (negative control), free TRAIL, or Exo-iNSC-TRAIL. At later time points, BLI shows some mice develop spinal tumors; **B**, tumor fold changes of *in vivo* MB231Br tumor growth at multiple time points after treatment by PBS, Free TRAL(C) and Exo-iNSC-TRAIL respectively, n = 6-7 mice per group, analyzed by two-way repeated measures ANOVA (\**P* < 0.05 and \*\**P* < 0.01); **C**, BLI quantification of individual tumor volumes normalized to tumor size at time of treatment; **D**, Kaplan-Meier survival curve for experiment described in (**A**)-(**C**), analyzed by Log-rank (Mantel-Cox) test (\*\**P*< 0.01). All data are shown as mean ± SEM.

The robust tumor killing effect of Exo-iNSC-TRAIL against MB231Br in vivo prompted us to test its efficacy in two additional GBM tumor models. Against the rapidly growing U87 line, a single dose of Exo-iNSC-TRAIL induced a statistically significant increase in survival compared to free TRAIL (p < 0.05) (**Fig. 7A-7C**). Against the invasive GBM8 glioblastoma line, three doses of Exo-iNSC-TRAIL also led to a statistically significant survival benefit compared to the untreated control (p < 0.01) and a trend toward significance when compared against free TRAIL (p < 0.07) (**Fig. 7D-7F**).

**Fig. 7.**
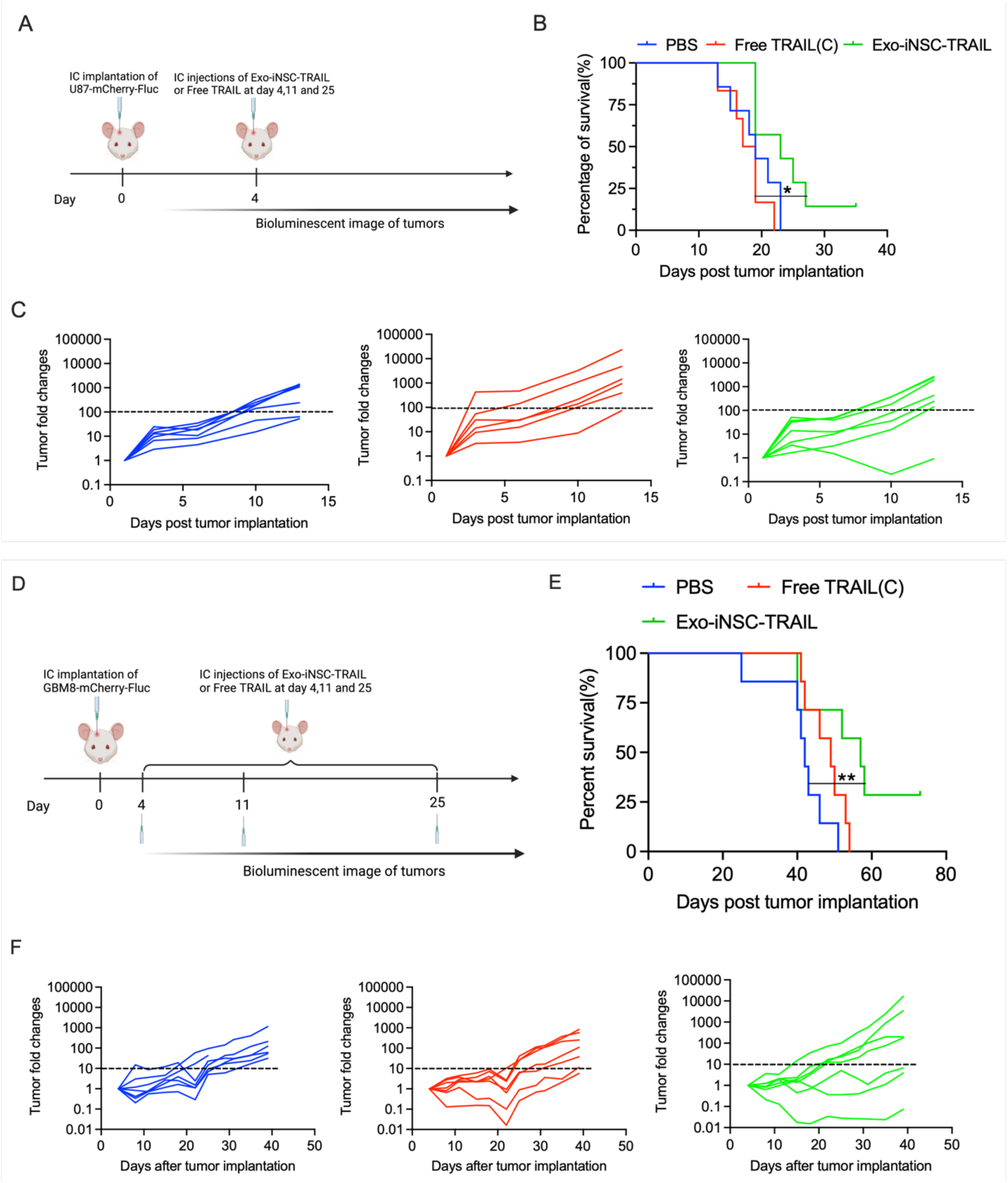
Measuring Exo-iNSC-TRAIL efficacy in two orthotopic *in vivo* models of glioblastoma. **A**, schematic of study design for GBM8 tumor treatment; **B**, Kaplan-Meier survival curve for mice with a brain intracerebral U87 tumor treated by PBS, Free TRAL(C) and Exo-iNSC-TRAIL respectively, n=6-7 mice per group, analyzed by Log-rank (Mantel-Cox) test (\**P* < 0.05);**C**, tumor fold changes of individual tumor volumes normalized to tumor size at day 1 after tumor implantation and treated by PBS, Free TRAL(C) and Exo-iNSC-TRAIL respectively; **D**, schematic of study design for GBM8 tumor treatment; **E**, Kaplan-Meier survival curve for mice with a brain intracerebral GBM8 tumor treated by PBS, Free TRAL(C) and Exo-iNSC-TRAIL respectively, n=7 mice per group, analyzed by Logrank test for trend (\*\**P* < 0.01); **F**, tumor fold changes of individual tumor volumes normalized to tumor size at the time of treatment by PBS, Free TRAL(C) and Exo-iNSC-TRAIL respectively. All data are shown as mean ± SEM.

## 4. Discussion

Here, for the first time, we identify EVs derived from engineered iNSCs as effective drug carriers for the pro-apoptotic agent TRAIL to improve its therapeutic efficacy in the treatment of brain cancers. We report that iNSCs auto-load EVs with a significant proportion of the TRAIL produced by those iNSCs, both within the lumen of the EVs as well as arrayed within the outer membrane. Exo-iNSC-TRAIL therapy significantly upregulated tumor apoptotic pathways and killing capacity against brain cancer cells *in vitro*, displayed selective accumulation in tumor cells *ex vivo*, and significantly increased survival in three orthotopic mouse models of human TNBC brain metastasis and GBM.

iNSCs were initially selected as the EV producer because they are rapidly proliferative in vitro, can be engineered to produce and secrete TRAIL, and display significant resistance to TRAIL-mediated apoptosis. Furthermore, iNSCs which secrete TRAIL are potent antitumor therapeutics themselves and contain cellular machinery which allows them to migrate through the brain, home to tumor foci, and deliver TRAIL to induce effective tumor killing [9],[7],[8]. Here, we identify, isolate, concentrate, and deliver the most potent form of TRAIL secreted by those iNSCs – Exo-iNSC-TRAIL – to maximize translational feasibility. Just as iNSCs display increased tumor-homing capacity compared to precursor NHF1 fibroblasts, Exo-iNSC-TRAIL contains a distinct proteome from Exo-NHF1-TRAIL, including greater TRAIL loading, which imparts increased tumor tropism *ex vivo*.

EVs have become a broadly used drug carrier to deliver various anti-cancer products with poor pharmacokinetics. Due to the poor bioavailability and low stability of the soluble recombinant TRAIL, the therapeutic efficacy of Dulanermin (a human recombinant TRAIL product from Amgen) in clinical studies is severely restricted even when employed at a high dose of as much as 30 mg/kg [51],[52],[53]. Thus, our initial desire was to load high levels of soluble TRAIL into the lumen of the EVs to decrease clearance, increase stability, and improve therapeutic efficacy. Interestingly, when we quantified the amount of TRAIL in EVs by ELISA, we found that we could also detect TRAIL on the surfaces of intact EVs; in fact, this surface-bound TRAIL comprised ∼75% of the total EV-associated TRAIL. Our immunoelectronmicroscopy and flow cytometry data both confirm that TRAIL was presented on the outer membrane of iNSC-derived EVs. Since the transfected TRAIL structure is soluble format and without the transmembrane domain of the natural TRAIL, the exact mechanism of EV membrane binding is not clear. One hypothesis is reflected in a recent paper showing that the Fas ligand (CD95L/Apo1L), which can be found both on the cell surface membrane and as a soluble protein and has a similar structure to TRAIL (aka Apo2L), can bind directly to fibronectin. Fibronectin is also found on EV surfaces, suggesting that this could be a membrane protein TRAIL can bind to in Exo-iNSC-TRAIL [54]. Furthermore, membrane-bound Fas ligand shows increased cytotoxicity compared to soluble Fas [55]. We observe a similar phenomenon in our studies: even *in vitro*, where no clearance occurs, Exo-iNSC-TRAIL is much more potent than free TRAIL (Fig 3C). This could be due to an enhanced multivalent interaction of TRAIL molecules on the surface of Exo-iNSC-TRAIL with DR4/5 on tumor cells, which could enhance DR4/5 clustering and in turn the tumor killing cascade. Exo-iNSC-TRAIL potency could be even further aided by the increased TRAIL stability afforded by association with the EV membrane (Fig 3E-G).

Previous studies have shown that EVs can stabilize TRAIL proteins and augment anti-cancer effects [56],[40]. TRAIL secretion via EVs has been identified as a naturally occurring method employed by various types of cells to regulate the activity and conduct of cells of interest in nearby or distant locations. For instance, human phytohemagglutinin (PHA)-activated T cells can produce TRAIL-anchored EVs to eliminate cells that are cancerous [57],[58]. Other cells like lymphoblast K562 cells and MSCs have also been engineered to make EVs, but these approaches either use cancer cells as EV producers or have very low yields of TRAIL loading [40],[56]. As an alternative method to improve TRAIL efficacy, mutation or multimerization of TRAIL to imitate native membrane-bound TRAIL has been investigated, such as the TRAIL-trimer fusion protein SCB-313 [59],[58]. TRAIL molecules arranged in such higher order oligomers may enhance DR5 clustering ability and in turn the tumor killing potency [59]. Moreover, when observed as a transmembrane protein, TRAIL has significantly more pro-apoptotic function than when produced as a soluble format, and this improved function is closely associated with its capability to assemble TRAIL receptors into supramolecular complexes [60],[56]. Together, these data support our findings that TRAIL incorporated into EVs has greater stability and an increased capacity to bind its appropriate receptor and stimulate the apoptotic cascade compared to free TRAIL [46],[48],[61].

The observed tumor tropism of iNSC-derived EVs is another important aspect of their potential as drug delivery vehicles. We have yet to determine whether this advantage is due to abundant surface-bound TRAIL acting as a targeting ligand to cells with high DR4/5, or whether other aspects of the unique proteome in Exo-iNSC-TRAIL play a role. Transdifferentiation of NHF1 cells into iNSCs imparts complex tumor-homing properties to this unique neural stem cell population, and while EVs are unable to “home” like living cells can, it is feasible that some relevant proteins could be transferred from iNSCs to Exo-iNSC-TRAIL which increase tumor tropism relative to Exo-NHF1-TRAIL. This phenomenon, and how it plays a role in the robust and sustained tumor growth inhibition *in vivo*, will be an important topic of future investigation.

To summarize, auto-loaded EVs derived from iNSCs are a promising and efficient treatment for brain cancer, particularly when used as local injection. Exo-iNSC-TRAIL can be easily produced, carries high levels of therapeutics, and improves anti-tumor efficacy compared to free TRAIL. This foundational study paves the way for a host of follow-up studies, including (1) testing Exo-iNSC-TRAIL against additional controls in vivo, including Exo-NHF1-TRAIL and the iNSC-TRAIL parent cells; (2) testing the pharmacokinetics, biodistribution, and therapeutic effect of Exo-iNSC-TRAIL after systemic intravenous injection; (3) engineering iNSCs to generate EVs loaded with other therapeutic proteins and/or additional tumor-targeting motifs, and (4) generating personalized EVs from iNSCs transdifferentiated from an individual patient’s own skin. Currently, little data exists regarding whether autologously generated exosomes for personalized treatment are superior to allogeneic, or off-the-shelf exosomes, and we are in a unique position to answer this question using exosomes from our transdifferentiated iNSCs.

### Statistical Analysis

All statistical tests and sample sizes are included in the Figure Legends. All data are shown as mean ± SEM. In all cases, the p values are represented as follows: ****p < 0.0001, ***p < 0.001, ** p < 0.01, *p < 0.05, and not statistically significant when p > 0.05. Mean values between two groups were compared using Student’s t-test or two-way ANOVA. Mean values between three or more groups were compared using one-way ANOVA. Mean values among three groups in longitudinal *in vivo* studies were compared using two-way repeated measurement ANOVA. All statistical analyses were performed using GraphPad Prism (Version 9.1.0). One outlying data point in Fig 3C was removed using Tukey’s Inter-Quartile Range method. For all quantifications of immunofluorescence, the samples being compared were processed in parallel and imaged using the same settings and laser power.

## ACKNOWLEDGMENTS

Dr. Satterlee would like to thank Professor Leaf Huang for his excellent mentorship and wishes him well in his retirement. This work was financially supported by the National Institutes of Health (R01NS099368); OBSC work was funded by the National Center for Advancing Translational Sciences grant U01TR003715. The UNC Proteomics Core Facility is supported in part by NCI Center Core Support Grant (2P30CA016086-45) to the UNC Lineberger Comprehensive Cancer Center. Some graphs were created by BioRender.com.

