## Supplementary material for "Auto-loaded TRAIL-exosomes derived from induced neural stem cells for brain cancer therapy": Zhang et al Supplementary Figures.pdf

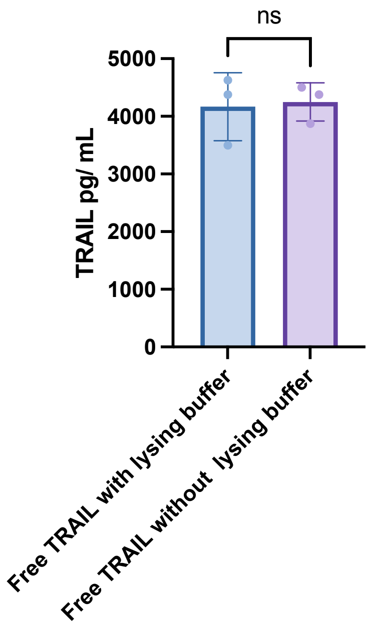


**Supplementary Fig. S1:** Comparison of the free TRAIL concentration by ELISA before and after addition of the lysing buffer + protease inhibitor

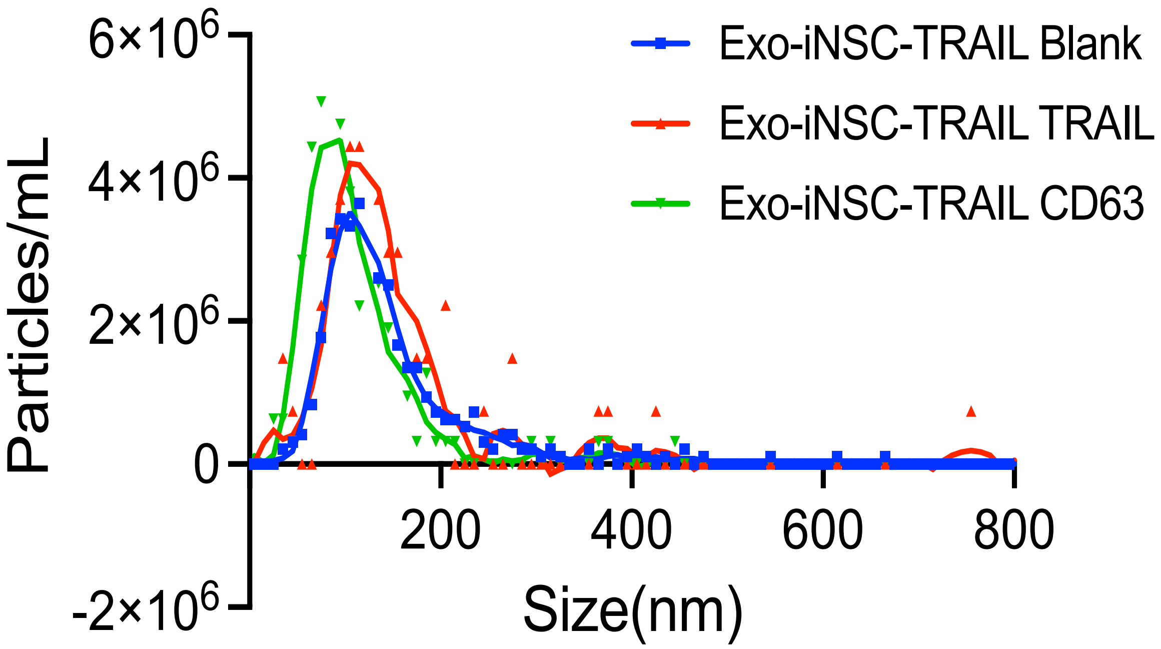


**Supplementary Fig. S2:** The size distribution of purified Exo-iNSC-TRAIL labeled with TRAIL or the exosomal marker tetraspanin (CD63)

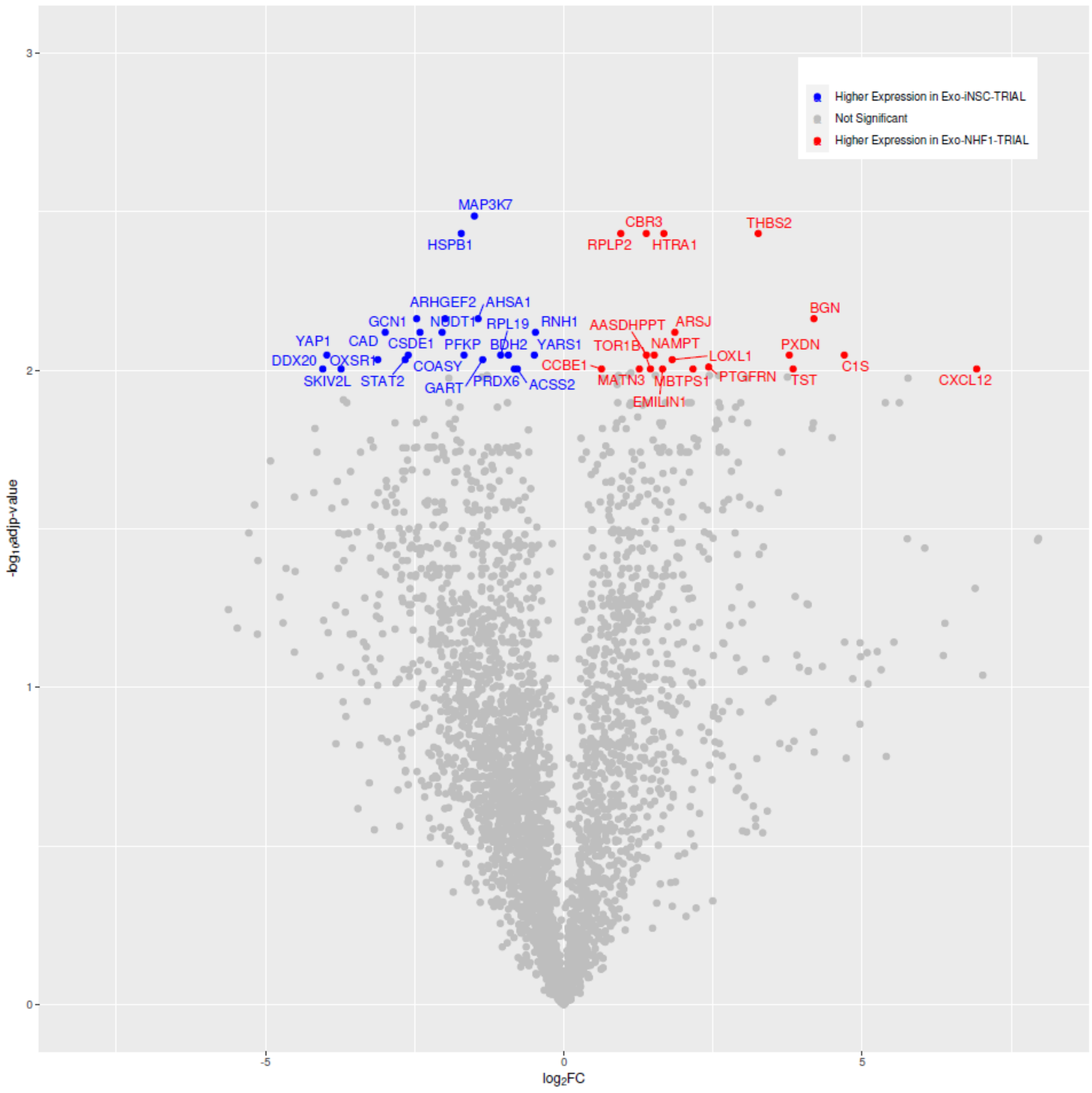


**Supplementary Fig. S3:** Volcano plot showing the results of the proteomic analysis of Exo-iNSC-TRAIL and Exo-NHF1-TRAIL (n = 3 biological replicates). Differentially abundant proteins with a FDR adjusted p-value < 0.01 are highlighted.

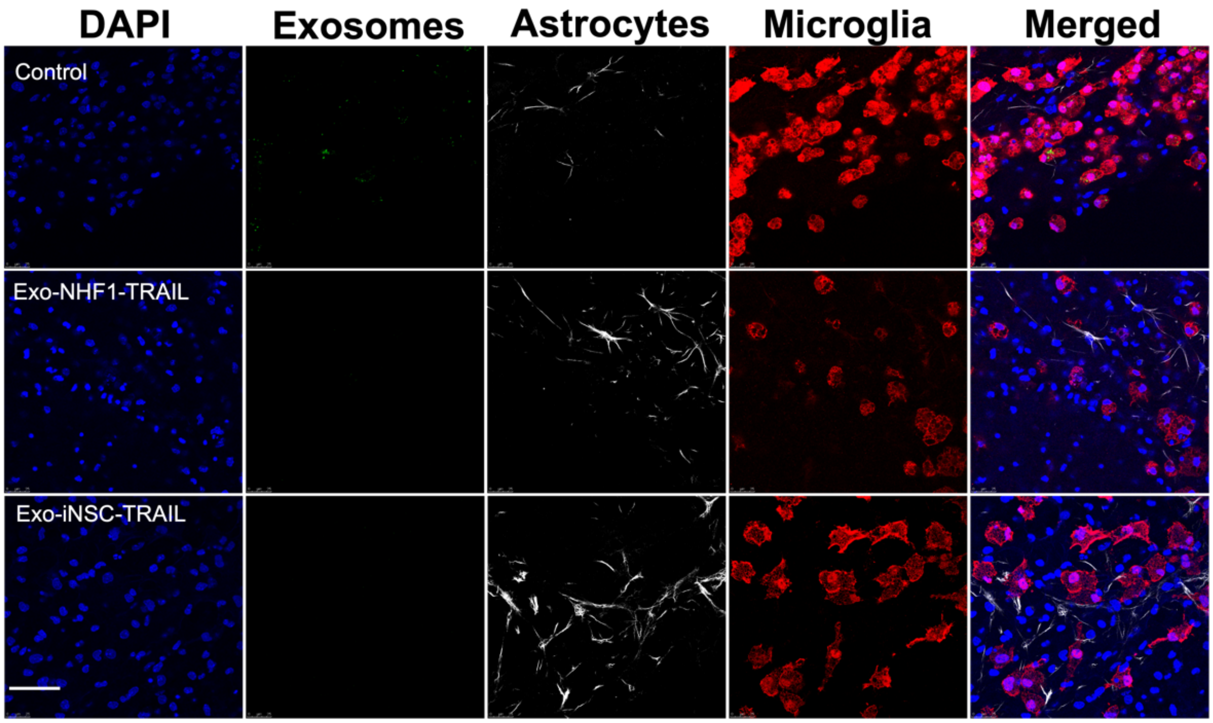


**Supplementary Fig. S4** Representative confocal images of fluorescently labelled Exo-NHF1-TRAIL and Exo-iNSC-TRAIL in non-tumor-bearing OBSCs. EVs were labelled by PKH26 (green), astrocytes were stained by GFAP (white), microglia were stained by IBA1+ (red), and nuclei were stained by DAPI (blue). t = 24h, scale bar = 50 μM.
